# Well-Lit: A programmable and customizable assistant for manual multi-well plate pipetting

**DOI:** 10.1101/2021.12.17.473010

**Authors:** Rafael Gómez-Sjöberg, Joana P. Cabrera, Andrew T. Cote

## Abstract

A very large number of biology and biochemistry laboratory protocols require transferring liquid aliquots from individual containers into individual wells of a multi-well plate, from plates to individual containers, or from one plate to another. Doing this by hand without errors, such as skipping wells, placing two samples in the same well, or swapping sample locations, especially when using plates with 96 wells or more, is difficult and requires enormous operator focus and/or a tedious manual error checking system. We present here a device built to facilitate error-free pipetting of samples from individual barcoded tubes to a multi-well plate or between multi-well plates (both 96 and 384 wells are supported). The device is programmable, modular and easily customizable to accommodate plates with different form-factors, and different protocols. The main components are only a 12.3” touch screen, a small form-factor PC, and a barcode scanner, combined with custom-made parts can be easily fabricated with a laser cutter and a hobby-grade 3D printer. The total cost is between approximately US$550 and US$600, depending on the configuration.

**Specifications table:** 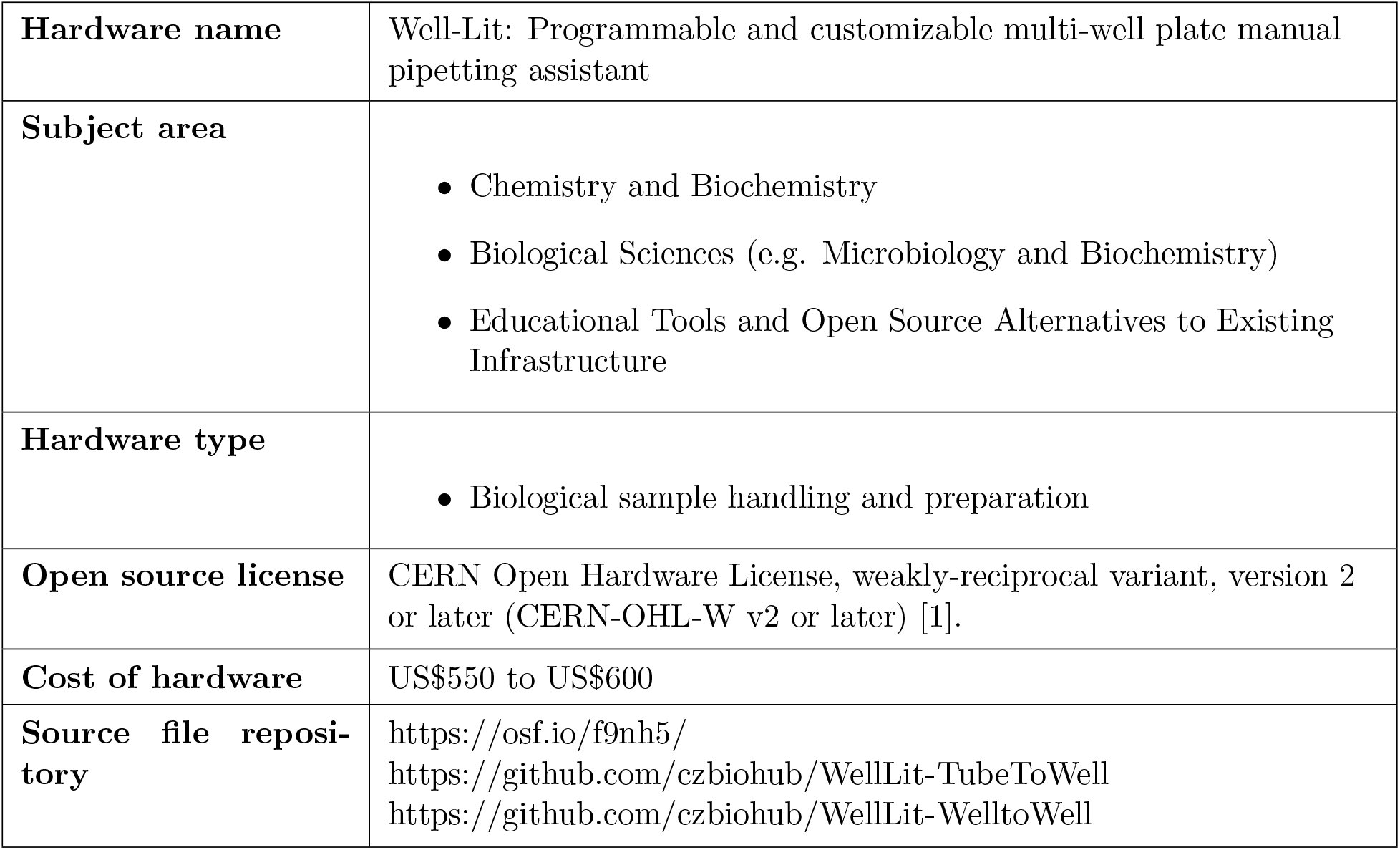

## 1. Hardware in context

At the beginning of March 2020, to address the dearth COVID-19 testing availability in the San Francisco Bay Area, the Chan Zuckerberg Biohub, in partnership with the University of California San Francisco Clinical Microbiology Lab (UCSF-CMBL), rapidly built and deployed an emergency COVID-19 viral testing facility [2]. The goal of this facility was to provide free, rapid, high capacity testing services to local hospitals, Departments of Public Health offices, and community-based clinical screening in the San Francisco Bay Area, with a particular emphasis on underserved populations. The facility was built in an extremely short time, returning the first clinical result 8 days after the start of the project.

Individual patient samples were arriving at the UCSF-CMBL as individual barcoded collection tubes. The testing protocol was run using multi-well plates, so at the start of the process the patient samples had to be transferred from each tube to an individual well on a barcoded 96-well plate, while keeping careful track of which patient sample goes to which well. This transfer had to be error-free, as any error would assign results to the wrong patient(s) or invalidate some or all results in a plate. Performing this transfer by hand without errors is a very slow and error-prone process. To minimize the chance of error, a manual process traditionally used at the UCSF-CMBL for the tube-to-well transfer involved two people, one doing the pipetting and the other registering the tube barcodes and writing down which well received each sample. Multiple steps of cross-checking between the two operators to avoid errors were necessary during the transfer, which increased the time needed to fill one 96-well plate to at least 2 hours. This manual process was too labor-intensive and slow, given that the testing facility needed to handle thousands of samples per day.

Several automated solutions for tube-to-plate transfer exist, but they are all based on liquid handling robotics that are usually very expensive, especially for high transfer throughput, and take a very long time to setup, program, and validate. Most liquid handling robots capable of performing this kind of transfer at the desired speed cost at least ~US$50,000. The testing facility purchased state-of-the-art liquid handling robots, but receiving them and getting them installed and validated for clinical deployment took several weeks. To immediately address the shortcomings of manual transfer, in the context of the pandemic emergency, the Biohub Bioengineering team designed and built, in only about 4 days, a simple device that facilitated an error-free manual tube-to-plate transfer process at the desired throughput.

After the initial deployment of the device in the UCSF-CMBL virus testing facility, it was also adopted by one other laboratory in the U.S. for the same COVID-19 testing purposes, also while their robotics systems were being set up. Additionally, scientists at the Biohub and at UCSF asked if the device could be adapted to consolidating cherry-picked wells from a set of source plates into a single destination well plate, for non-COVID-19 related projects. Thanks to the size of the screen, and the fact that the initial design was modular, it was very easy to create a new configuration of the device to perform this well-to-well transfer. We call the basic framework of the device “Well-Lit”, and the two configurations are called “Tube to Well-Lit” (sample tube to well-plate transfers), and “Well-Lit to Well-Lit” (well-plate to well-plate transfers). We provide designs for holders for four different brands and types of well plates, which can be used with either configuration, and it is very easy to design and 3D print new holder designs, and to adapt the device to completely new protocols.

At least two open-source devices that perform a similar function have been described [3, 4], but they both use an array of LEDs (light-emitting diodes) to illuminate the wells from the bottom. This design has two important disadvantages compared to our LCD screen-based design: (1) It requires the construction of custom electronic circuitry, which many labs do not have the expertise and equipment to do, and (2) the position and color of the illumination pattern are fixed so they cannot be reprogrammed to work with different types of well plates or protocols, or to make the device usable by color-blind individuals. Gilson Inc. (Middleton, WI) sells the “TRACKMAN Connected” system [5], which functions very similarly to our device (using a tablet instead of a touch screen), but it costs about three times more and is designed to function exclusively with Gilson-brand pipettes - it is a “closed-source” system that cannot be easily modified by the user. Additionally, the Gilson device can only accept a single plate at a time, so it cannot help with the transfer of samples from one plate to another. The biggest advantage of our open source design is that it can be easily reprogrammed to suit the specific needs of the user, such as changing the colors of the wells for color-blind users, accommodating non-standard well plates, adding complex custom protocols, etc.

## 2. Hardware description

We named this device “Well-Lit” because of the way it functions. The multi-well plates rest on top of a computer touch-screen, held in place by a custom-made holder, and an array of brightly colored discs is displayed on the screen underneath the plate, one disc per well, so that the wells are illuminated with different colors to indicate to the operator which wells already have samples, and which well should receive the next sample (Figure 1). In the “Tube to Well-Lit” configuration, each sample tube is read by the barcode scanner, which triggers the software to illuminate the next available well in the receiving plate. Due to the extreme time pressure, the device design was necessarily based almost exclusively on parts already in-hand. We had no time to shop for a different screen, for example. The device is composed of a 12.3” computer touch-screen (scavenged from an older project), a miniature Windows PC computer (removed from a screen used to display conference room occupancy), a barcode scanner (for scanning the plate and sample tube barcodes), and various custom-designed parts that are laser-cut and 3D-printed in-house.

**Figure 1:**
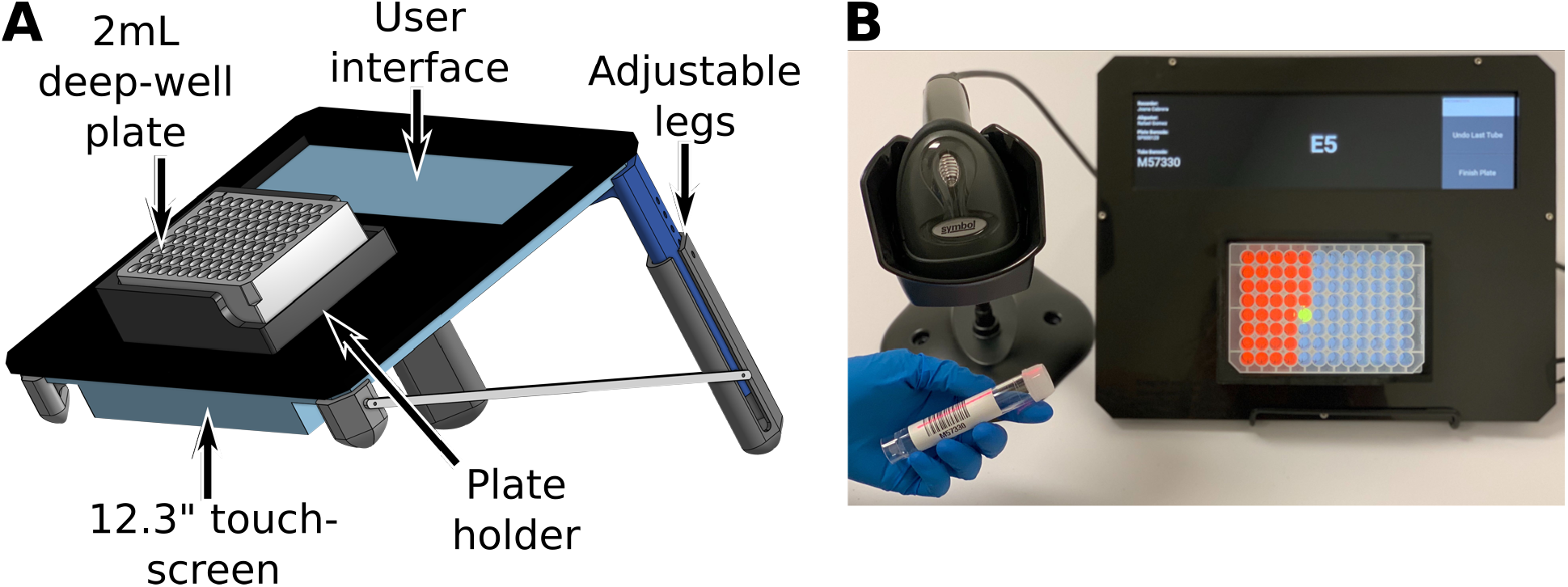
Well-Lit device developed to expedite the manual tube-to-well pipetting process (“Tube to Well-Lit” version), while simultaneously minimizing the chance for error. (A) CAD rendering showing the basic structure of the device: 3D printed legs hold a 12.3” touch-screen at an angle to allow the operator to see straight into the wells in the plate (standard legs are designed to allow adjusting the angle between approximately 30 and 46 degrees with respect to the table). A plastic panel with two laser-cut openings, one for the multi-well plate, and the other for a user interface area, is placed over the screen. A 3D printed plate holder bolted to the panel provides a key so that the plate can only be inserted in one orientation, and also shades the sides of the plate from ambient light to improve the contrast of the color discs illuminating the wells. The front panel is designed to be easy to place and remove for decontamination, and to allow panels designed for different types of multi-well plates to be easily swapped by the operator. (B) The Well-Lit in operation, with the barcode scanner used to scan the barcodes on the multi-well plate and patient sample tubes. Wells that have already received samples are illuminated in red, while the well that should receive the sample from the tube that was just scanned is illuminated in yellow. The user interface area shows the operator name, plate barcode, sample barcode, and current well. Buttons on the right end of the interface allow the user to undo the last tube scan and finish the plate.

Use of the “Well-Lit” device drastically reduces the chance of error by providing clear visual guides for each sample, while simultaneously tracking and checking input barcodes. During each transfer run the device generates a Comma-Separated Values (CSV) file recording all the transfers that were performed.

Thanks to the modular design, the plate holder can be easily re-designed to accept different types of plates, and the software can be easily modified to handle different protocols. Here we present two versions of the device: The standard configuration described so far and shown in Figure 1 (“Tube to Well-Lit”), and a version designed to help cherry-pick samples from one or more source multi-well plates to a single destination multi-well plate (“Well-Lit to Well-Lit”) (Figure 2). The software in this second version reads a CSV file specifying a set of source plates and wells in those plates with the corresponding destination wells in the destination plate.

**Figure 2:**
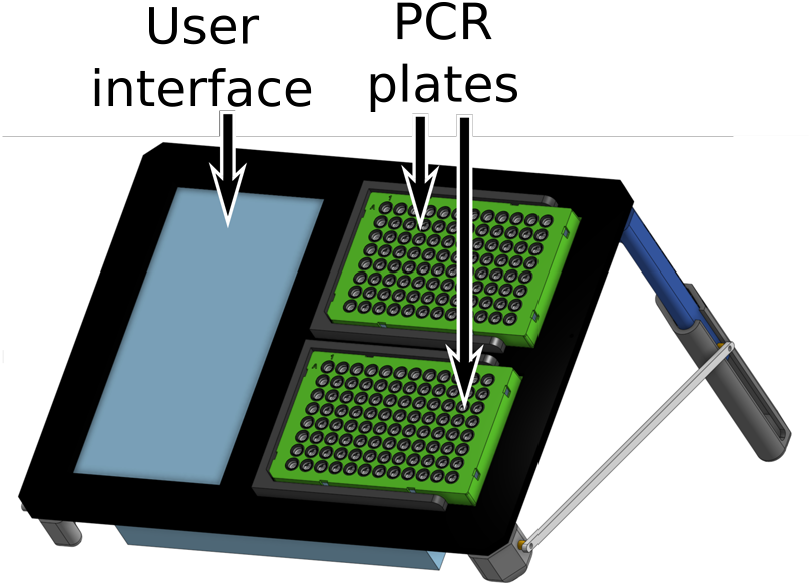
“Well-Lit to Well-Lit” version of the device, developed to expedite the manual cherry-picking of aliquots from one or more 96- or 384-well PCR plates to a single plate, or re-mapping the samples from one plate to another, while simultaneously minimizing the chance for error. This version does not need a barcode scanner. The software in this version reads a CSV file specifying the source plate and well and destination well for each pipetting step.

We provide holder designs for Thermo Fisher 2 mL 96-well plates (cat# 12-566-612), Bio-Rad low profile 96-well PCR plates (cat# HSP-9641), USA Scientific 1 and 2 mL 96-well plates (cat# 1896-1000 & 1896-2000), and Thermo Fisher Matrix 96-tube racks (cat# 4897-BR), that can be used with either version of the device. It is important to note that the only thing needed to switch the device between “Tube to Well-Lit” and “Well-Lit to Well-Lit” configurations is a different front panel, and running a different version of the software, so a single unit can serve both roles very easily.

For both devices we provide two adjustable screen leg designs, the standard one that allows the screen angle to be adjusted between approximately 30° and 46° with respect to the table, and one that provides an angle range between approximately 20° and 33°. This second set of legs is best used when the liquid level in the multi-well plates (or Matrix tubes) is high, and steeper angles would lead to spills. The device can be used without the legs, with the screen lying flat on the bench, at the cost of decreased visibility of the color patterns under the wells, unless the user can look straight down into the wells during the pipetting operations.

To summarize, this device is most helpful for minimizing the chance of error when performing the following procedures by hand:

- Transfer of aliquots from individual containers to a multi-well plate, with the aliquot from each container going to a single well.
- Cherry picking aliquots from one or more source multi-well plates into a single destination multi-well plate.
- Re-mapping (re-arraying) the samples from one multi-well plate to another.

The programmability and modularity of the device make it very easy to adapt to many other uses.

## 3. Design files

### 3.1 Design Files Summary

All design files are made available under the CERN Open Hardware License version 2 or later, weakly-reciprocal variant (CERN-OHL-W v2 or later) [1].

- **Well-Lit** - This is not strictly a file, but a document in a cloud-based CAD system called Onshape (onshape.com), available for free for non-commercial use. This document has the full 3D model of the device and all of its components, allowing anybody to explore all the details of the design of the device, and to export all the STL and DXF files listed in Table Additionally, it is possible to download the components and assemblies as STEP, IGS, or other standard 3D file formats. The document can also be copied to a user’s account to make modifications and create new versions of it.
- **WellLit-TubeToWell** - This repository contains the software needed to run the device in the Tube to Well-Lit configuration. Note: This repository is automatically linked to a common Github repository called WellLit.
- **WellLit-WellToWell** - This repository contains the software needed to run the device in the Well-Lit to Well-Lit configuration. Note: This repository is automatically linked to a common Github repository called WellLit.

## 4. Bill of materials

### 4.1 Main Off-the-shelf components

Table 2 contains the list of major off-the-shelf components. Note that only one of each component is needed. A few things to keep in mind about these components:

**Table 1:**
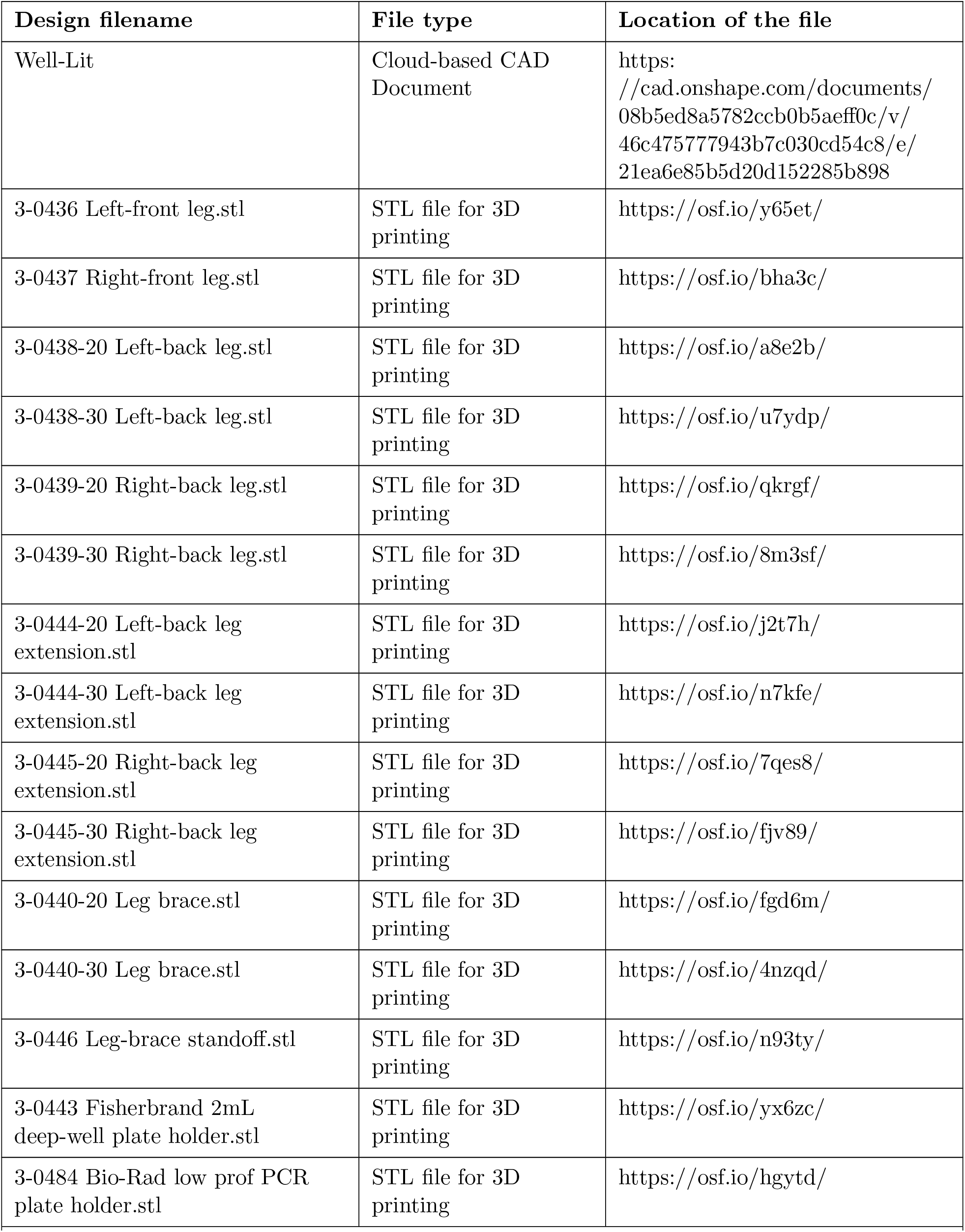
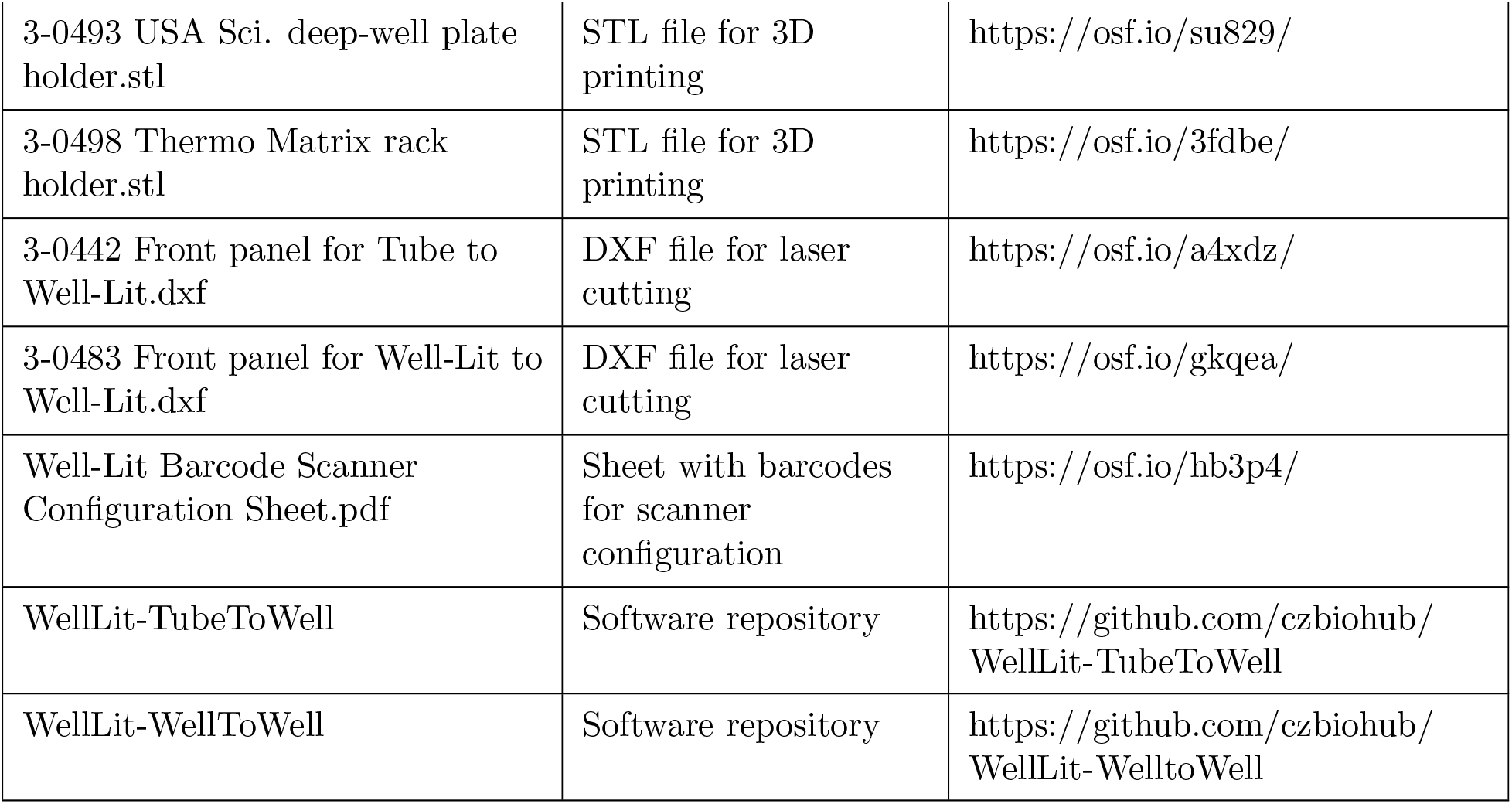
Design files summary.

**Table 2:** Main off-the-shelf components

| Designator | Description | Cost<br>US\$ | Source of materials |
| --- | --- | --- | --- |
| Screen | UPERFECT 12.3" touch-screen | 179.99 | https://www.amazon.com/gp/product/B07YWZH3TZ |
| PC | Mini PC with Windows 10 Pro | 149.99 | https://www.amazon.com/gp/product/B07QY8LDGX |
| Keyboard | Logitech compact Bluetooth keyboard with touch pad | 59.99 | https://www.amazon.com/gp/product/B00ZOPVSKW |
| Scanner | Symbol DS4308 barcode scanner with stand | 153.00 | https://www.amazon.com/gp/product/B0799R5MXS |

- A different screen of similar size could be used instead of the one specified above, but the legs and front panel would have to be redesigned.
- Any small form-factor PC with Windows will work, as long as it has one HDMI port, and at least 3 USB ports. We chose a Windows computer because it was very easy to integrate into the existing IT infrastructure in our labs. A PC running Linux, MacOS, or even a Raspberry Pi could be used with minor modifications to the software.
- Any other keyboard will work - we like the one specified in the table because it is very small and has an integrated touch pad, which are important features when trying to use the device inside a small and/or crowded biosafety cabinet.

### 4.2 Minor off-the-shelf components

The minor off-the-shelf components listed in Table 3 are not sold in the quantities needed to build one device from the source indicated on the table. Screws and nuts are sold in packs (usually 25 to 100 per pack), and obviously tape is sold in a roll. The per-unit costs indicated in the table were calculated from the cost of a pack or roll from the indicated source. Buying individual screws and nuts from a local store will be more expensive. We used hot glue instead of epoxy glue, because hot glue is much easier to remove in case the build requires fixing or modifications, but epoxy glue will provide a much stronger and more durable bond.

**Table 3:** Minor off-the-shelf components. The number of plate holder screws depends on the type of device being built. The “Tube to Well-Lit” needs only 8 for a single plate holder, while the “Well-Lit to Well-Lit” requires 12, since it has two plates.

| Designator | Description | Qty | Unit Cost US\$ | Total Cost US\$ | Source of materials |
| --- | --- | --- | --- | --- | --- |
| Leg brace screws | Thread-forming screws for plastic, rounded head, stainless steel, #2, 19 mm (3/4”) long, cat# 96001A164 (can substitute with M3, 20 mm long screws) | 4 | 0.22 | 0.91 | McMaster-Carr |
| Plate holder screws | Thread-forming screws for plastic, flat head, stainless steel, #2, 12.7 mm (1/2”) long, cat# 95893A560 (can substitute with M3, 15 mm long screws) | 6 or 12 | 0.27 | 1.61 or 3.22 | McMaster-Carr |
| Double-sided tape | Very-High-Bond (VHB) Foam Mounting Tape for Hard-to-Bond Materials, 19 mm (3/4”) wide, cat# 76675A33 | 165 mm (6.5”) | 0.27 | 1.75 | McMaster-Carr |
| Panel screws | Socket-head screws, stainless steel, #4-40, 12.7 mm (1/2”) long, cat# 92196A110 (can substitute with M3, 15 mm long screws) | 5 | 0.04 | 0.21 | McMaster-Carr |
| Nylon nuts | Nylon hex nut, #8-32, cat# 94812A400 (can substitute with M4 nut) | 4 | 0.07 | 0.30 | McMaster-Carr |
| Nylon screws | Nylon screw, #8-32, 1.5” long, cat# 94735A743 (can substitute with M4 screw) | 2 | 0.08 | 0.16 | McMaster-Carr |
| Glue | 2-part or UV epoxy adhesive |  |  |  | Any |
| Silicone sealant | General purpose silicone sealant |  |  |  | Any |
| Masking tape | General purpose masking or painter’s tape |  |  |  | Any |

### 4.3 Custom-made components

The Well-Lit has two main sets of custom-made components, one set is for the legs that support the screen (see Figure 3, which also shows purchased components needed for the leg assembly), and the other for the front panel that holds the multi-well plate over the screen (Figure 4). All custom-made components are either 3D printed, or laser-cut. Table 4 shows all of the components that should be 3D printed. Any 3D printer of the right size can be used to print all of these components. The back legs and back leg extensions come in two varieties, depending on the screen angle range desired. Their part numbers have a “-20” suffix for the 20°-33° range, and “-30” for the 30°-46° range. Choose only one angle for a particular build, as parts designed for different angles cannot be mixed together or easily swapped after the device is built.

**Table 4:** 3D printed components. The back legs and back leg extensions come in two varieties, depending on the screen angle range desired 20°-33° or 30°-46°. Their part numbers have a “-20” or “-30” suffix, depending on the angle range. The number of plate holders to print depends on the type of device being built (“Tube to Well-Lit” or “Well-Lit to Well-Lit”) and the type of plate used for it. Any plate type can be used in either version, and two different plates types can be used in the “Well-Lit to Well-Lit”.

| Designator | Description | Qty. | Unit Cost (US\$) | Total Cost (US\$) |
| --- | --- | --- | --- | --- |
| 3-0436 | Left-front leg for screen | 1 | \$0.36 | \$0.36 |
| 3-0437 | Right-front leg for screen | 1 | \$0.36 | \$0.36 |
| 3-0438-20 or 3-0438-30 | Left-back leg for screen | 1 | \$1.44 | \$1.44 |
| 3-0439-20 or 3-0439-30 | Right-back leg for screen | 1 | \$1.44 | \$1.44 |
| 3-0444-20 or 3-0444-30 | Left-back leg extension for screen | 1 | \$1.56 | \$1.56 |
| 3-0445-20 or 3-0445-30 | Right-back leg extension for screen | 1 | \$1.56 | \$1.56 |
| 3-0440-20 or 3-0440-30 | Leg brace | 2 | \$0.20 | \$0.40 |
| 3-0446 | Standoff for leg braces | 4 | <\$0.01 | <\$0.04 |
| 3-0443 | Holder for 2 mL deep-well plate (Fisher Sci. cat# 12-566-612) | 0, 1 or 2 | \$2.16 | |
| 3-0498 | Holder for Thermo Matrix tube rack | 0, 1 or 2 | \$1.60 | |
| 3-0484 | Holder for Bio-Rad low profile PCR plate (Bio-Rad cat# HSP-9641) | 0, 1, or 2 | \$0.56 | |
| 3-0493 | Holder for USA Scientific 1 and 2 mL deep well plates (USA Sci. cat# 1896-1000 & 1896-2000) | 0, 1, or 2 | \$2.20 | |

**Figure 3:**
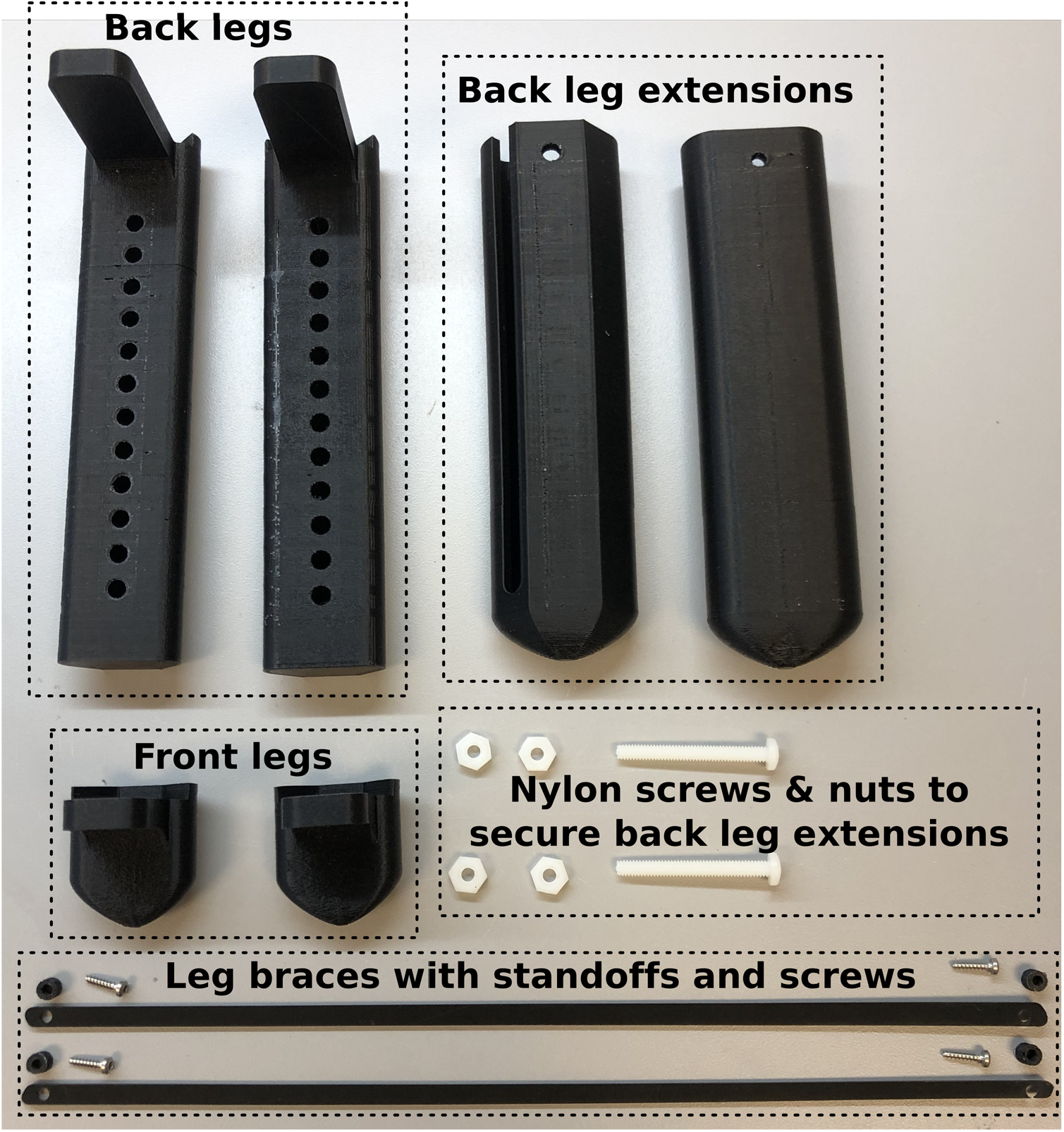
Components for the legs that hold the screen. The legs, leg extensions, and leg brace standoffs are all 3D printed. The leg braces are laser-cut.

**Figure 4:**
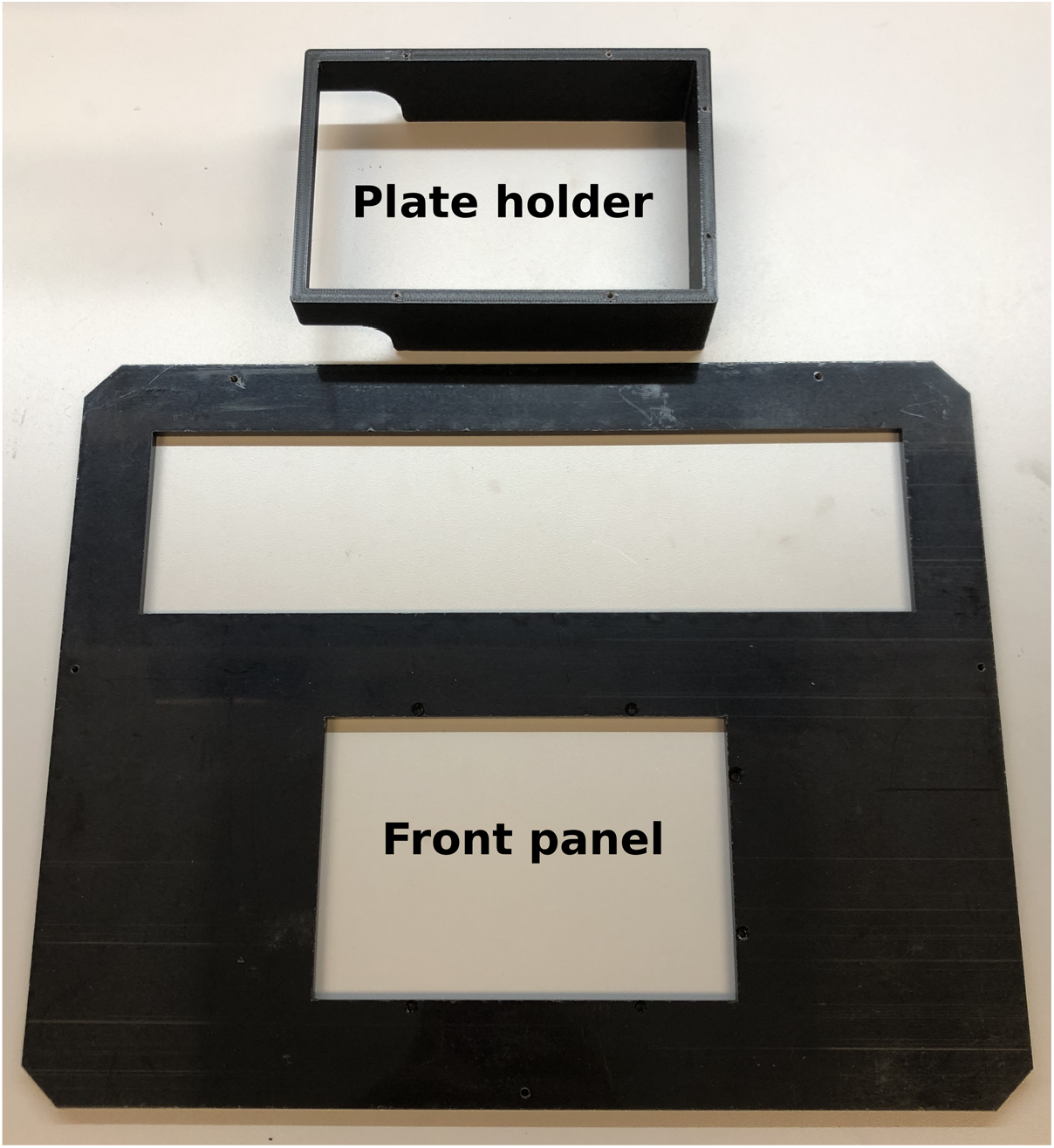
Components for the “Tube to Well-Lit” front panel, showing the 3-0443 plate holder. The plate holder is 3D printed. The panel is laser-cut.

Table 5 lists all the parts that should be laser-cut. It is strongly recommended that all lasercut parts be made from acetal sheets (instead of acrylic) because acetal is much more resistant to the cleaning agents typically used to sanitize the device after use, such as ethanol or bleach solutions. Additionally, acetal is a lot less brittle than acrylic, which improves the longevity of the parts. However, if acetal is difficult to find, acrylic would work at the cost of reduced longevity. The thickness of the acetal sheets used for laser-cutting is not particularly critical, especially when converting from imperial to metric thicknesses, so the closest available thickness can be used without trouble. The cost of the laser-cut parts was estimated based on the price of a 4.76 mm (3/16”) thick sheet of acetal 304.8 mm (12”) by 609.6 mm (24”) in size.

**Table 5:**
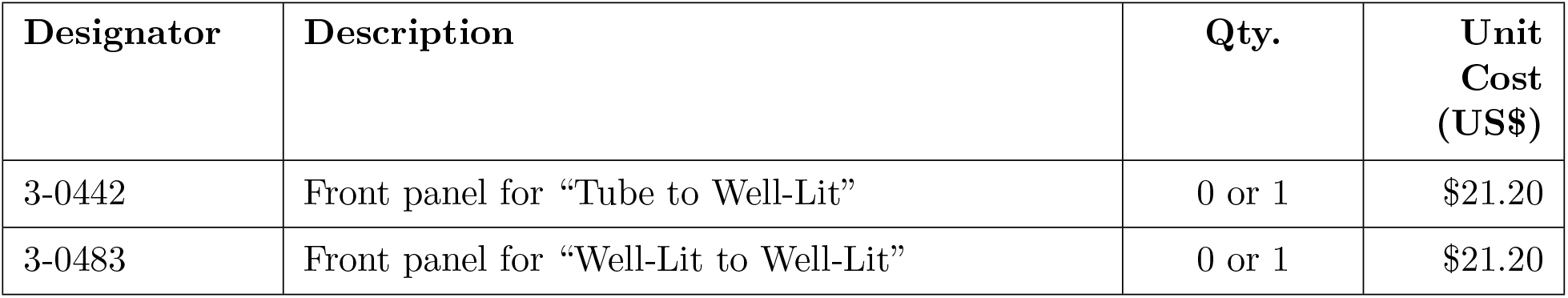
Laser-cut components. All of them are cut from 4 to 5 mm (3/16”) thick acetal sheets. The number of front panels depends on the type of device being built (“Tube to Well-Lit” or “Well-Lit to Well-Lit”).

The screen-plus-legs assembly is common to both versions of the device (“Tube to Well-Lit” and “Well-Lit to Well-Lit”). The only thing that is different between the two versions is the front panel, which is removable/swappable, and the software. Front panels for multiple versions of the device and for different types of plates can be built to be able to use a single screen for a variety of protocols. The number of plate holders to print depends on the version of device being built. The “Tube to Well-Lit” version only has one plate holder on the front panel, while the “Well-Lit to Well-Lit” version has two holders. Any plate type can be used in either version, and two different plates types can be used in the “Well-Lit to Well-Lit”.

Important: The STL and DXF files for the printed and laser-cut parts are all in millimeters (1 unit = 1 mm).

## 5. Build instructions

### 5.1 Screen legs

1. Parts required:
  - Screen
  - Front legs (3-0436, 3-0437) - 3D printed
  - Back legs (3-0438, 3-0439) - 3D printed
  - Leg braces (3-0440) - 3D printed
  - Leg brace standoffs (3-0446) - 3D printed
  - Double-sided tape
  - Glue
  - Nylon screws
  - Nylon nuts
  - Leg brace screws
  - Masking tape
  - Silicone sealant
2. If using M4 nylon screws (instead of #8-32), the large holes on the back legs might need to be drilled to provide the right clearance for an M4 screw.
3. Put double-sided tape on the ledges of all legs, as shown in Figure 5. The tape will secure the legs to the back of the screen.
4. Mount the legs onto the back of the screen as shown in Figure 6, aligning each leg square with its respective corner. Make sure that you match the left and right legs properly, as shown in Figure 6A. Each leg has a small screw hole on the side for attaching a leg brace. Make sure you mount each set of legs with the brace screw holes facing the same direction, always away from the screen. Press all the legs firmly down to ensure that the tape is well adhered.
5. Apply a bead of glue all around the ledge of each leg as shown in Figure 6C. This glue is very important to make the legs sturdy, as the double-sided tape is not strong enough by itself.
6. Mount a leg brace on each set of legs (left and right) using the leg brace screws (rounded head thread-forming screws), putting the brace standoffs between the legs and the braces, as shown in Figure 7.
7. Tap the hole in the wide flat face of each leg extension with a #8-32 tap (or M4). As indicated in the figure below, the hole on the narrow flat face of the extension has clearance for the screw (it might have to be drilled for proper clearance if using M4 screws). Insert the tap through the clearance hole, as shown in Figure 8, to keep the threads well aligned with the clearance hole.
8. Put two nylon nuts into each nylon screw (#8-32 or M4) and tighten each nut firmly against the head of the screw, as shown in Figure 9. These nuts make it easier to grab and turn the screws by hand when using the device.
9. Slide each leg extension over its respective back leg, and set their position with the nylon screws, as shown in Figure 10. The extensions are used to set the best angle for the screen, depending on the height of the surface where it is used, the type of plate, and the height of the user.
10. Cover the front bezel of the screen with masking tape, with the tape flush against the edge of the bezel that goes down to the screen, and make a frame with mask tape directly on the screen, leaving a 1-2 mm gap between the tape and all the edges of the bezel. See Figure 11A.
11. Apply a bead of silicone sealant all around the inner edge of the screen (into the 1-2 mm gap between the tape frame and the bezel) to create a water-proof seal between the edge of the screen and the bezel. See Figure 11B. Let the silicone cure fully, according to the manufacturer instructions. This seal extends the life of the screen by preventing liquids, from accidental spills, or from spraying the device with sanitizing solutions, from seeping into the electronics through the gap between the screen and the bezel.
12. Remove the masking tape from the bezel and the screen. Using a sharp x-acto knife to cut the silicone along the edges of the tape on the screen (not on the bezel) might make the tape easier to remove without lifting the silicone seal.

**Figure 5:**
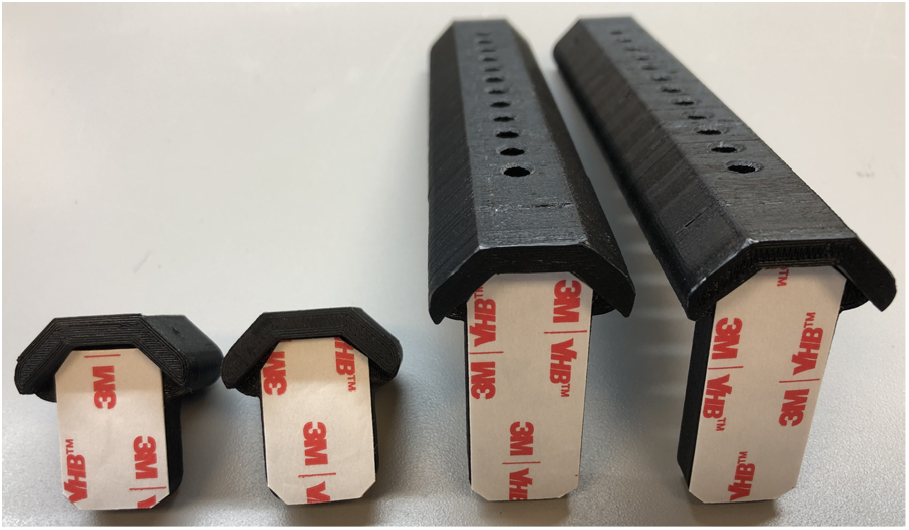
Double-sided tape applied onto the ledges of all the screen legs.

**Figure 6:**
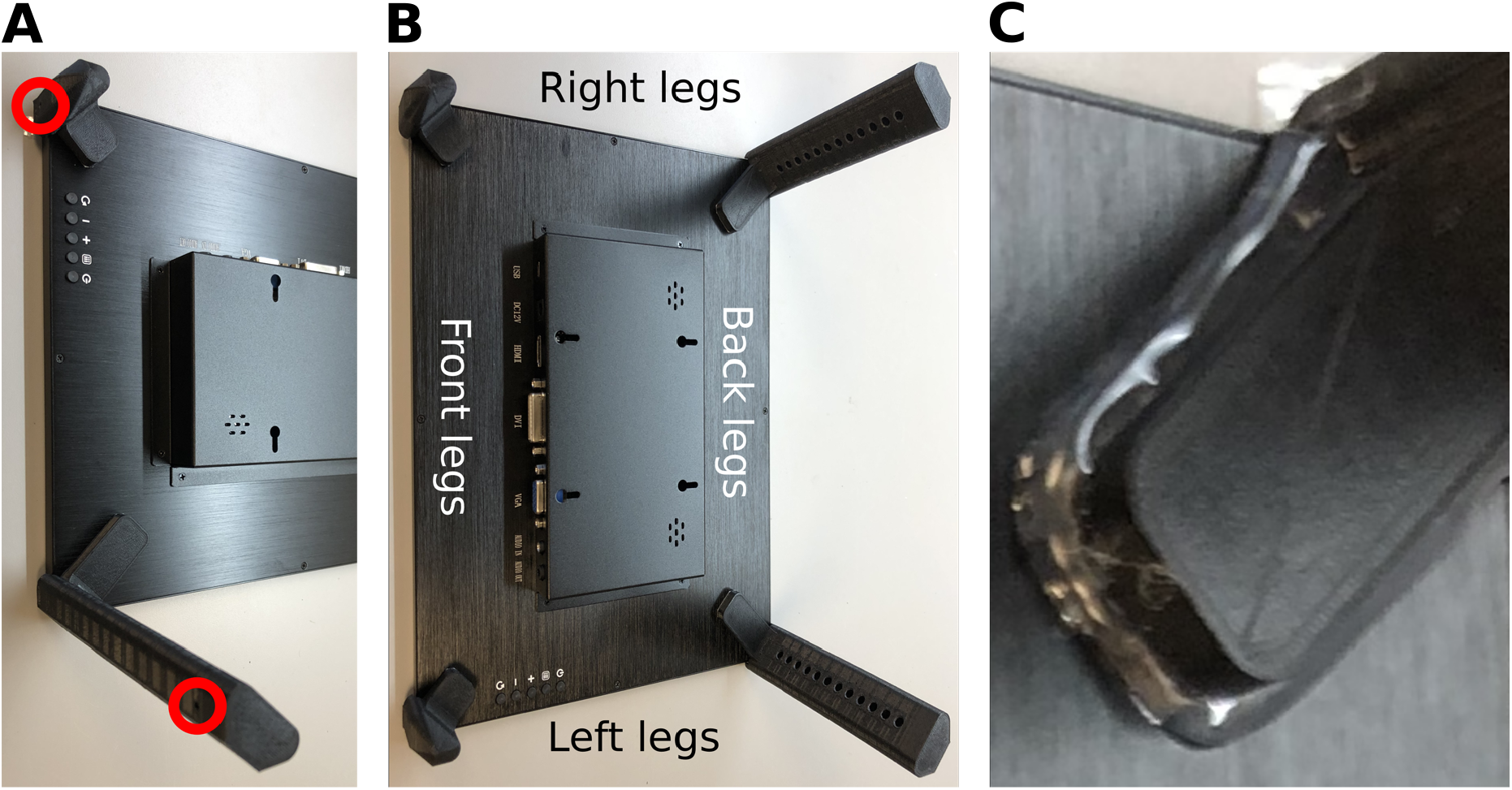
Legs attached to the back of the screen. (A) Each set of legs (left and right) have matching holes for mounting the leg braces (highlighted by the red circles) that should face away from the screen. (B) Notice the location and orientation of all legs with respect to the buttons and connectors on the screen. (C) Glue applied all around the ledge of a leg.

**Figure 7:**
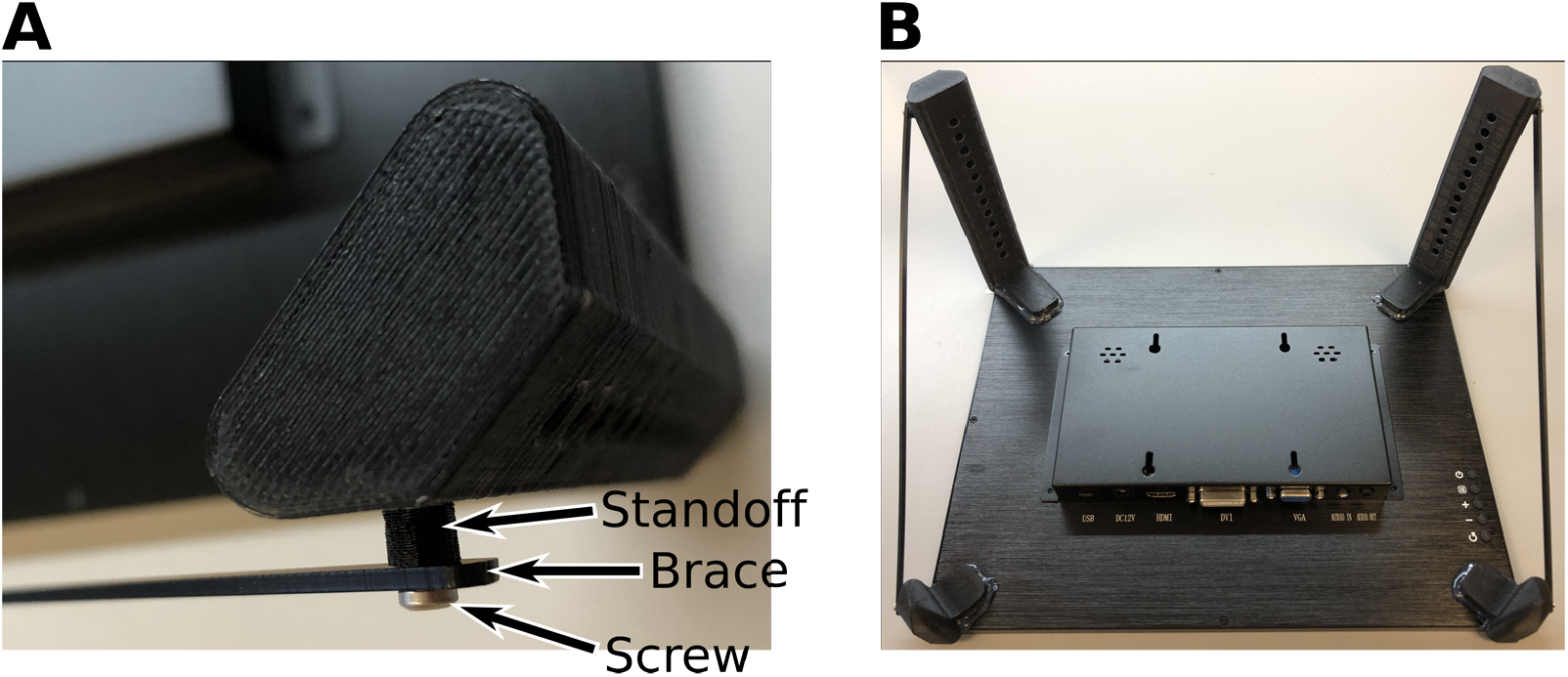
Braces attached to the legs. (A) Notice the order in which the brace and standoff are placed. (B) Both braces, left and right, attached to the legs.

**Figure 8:**
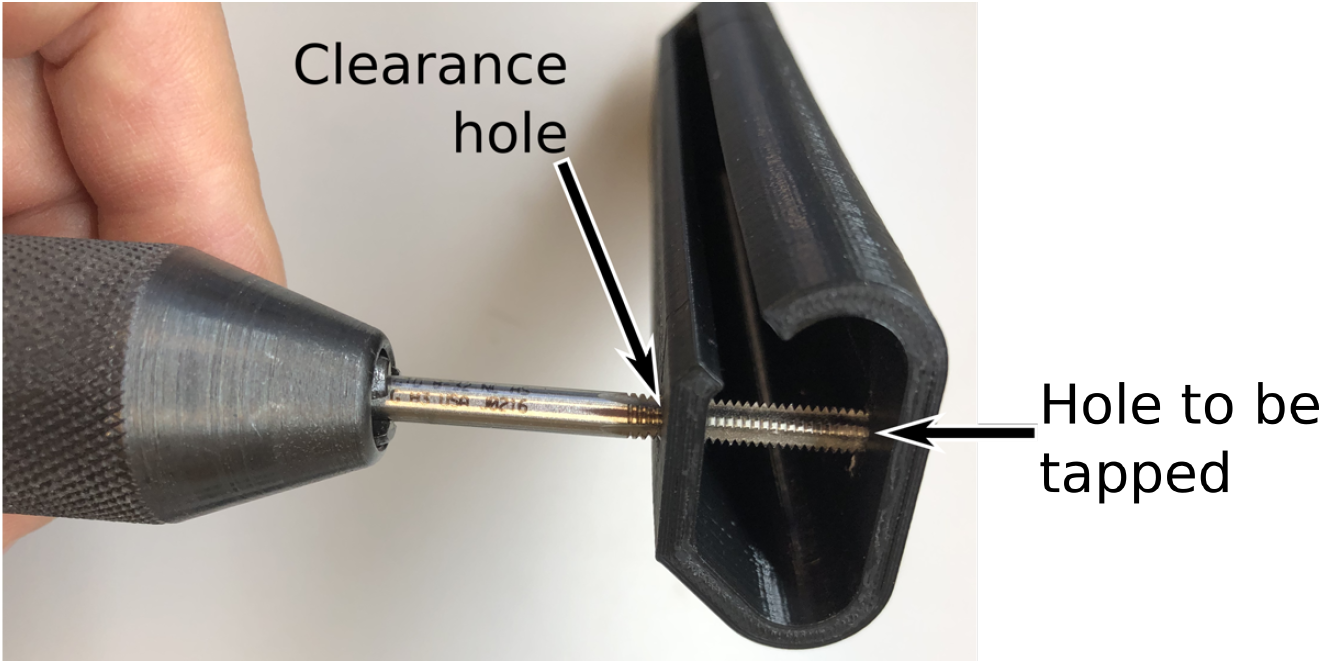
Tapping the hole on the wide flat side of the leg extensions.

**Figure 9:**
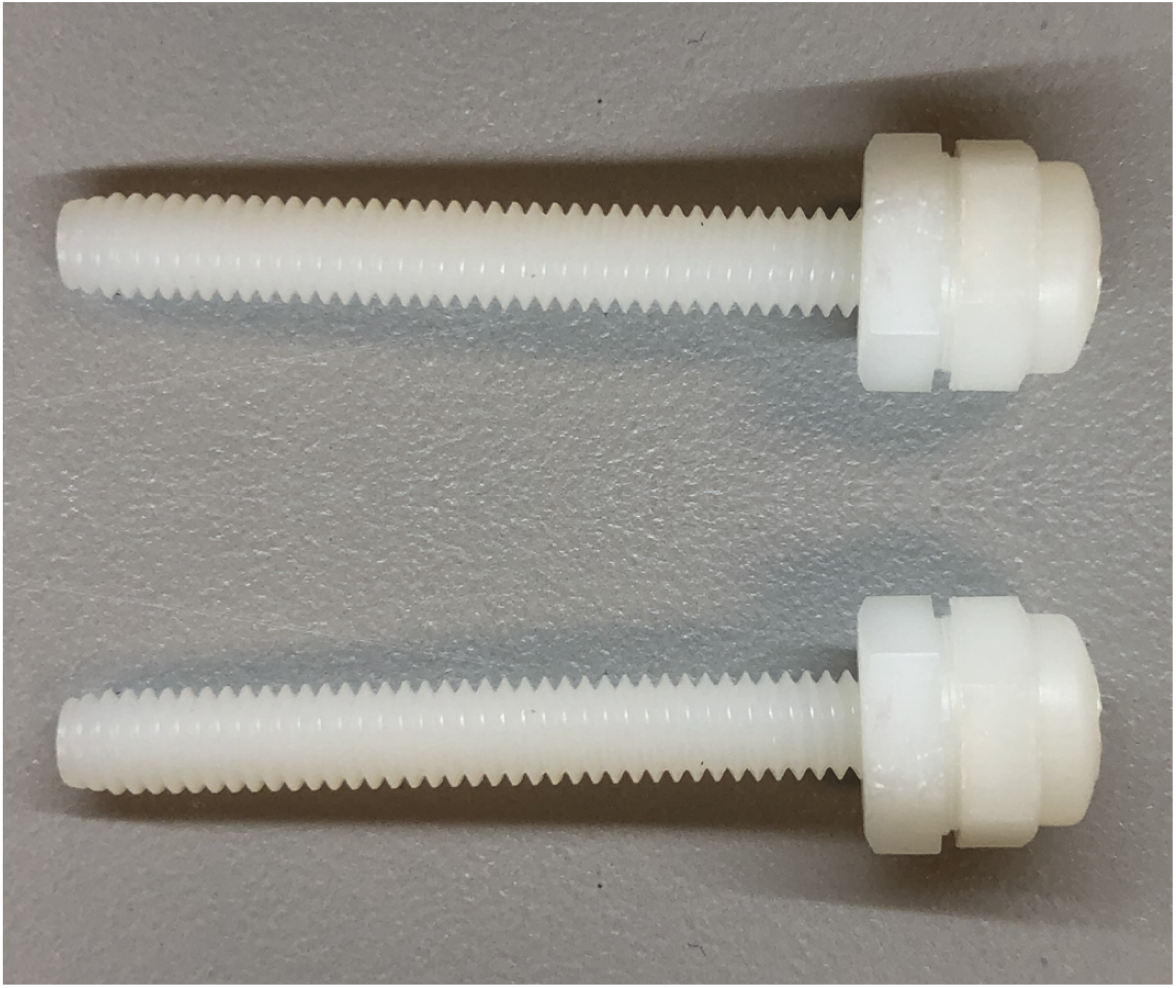
Nylon nuts tightened against the head of the nylon screws.

**Figure 10:**
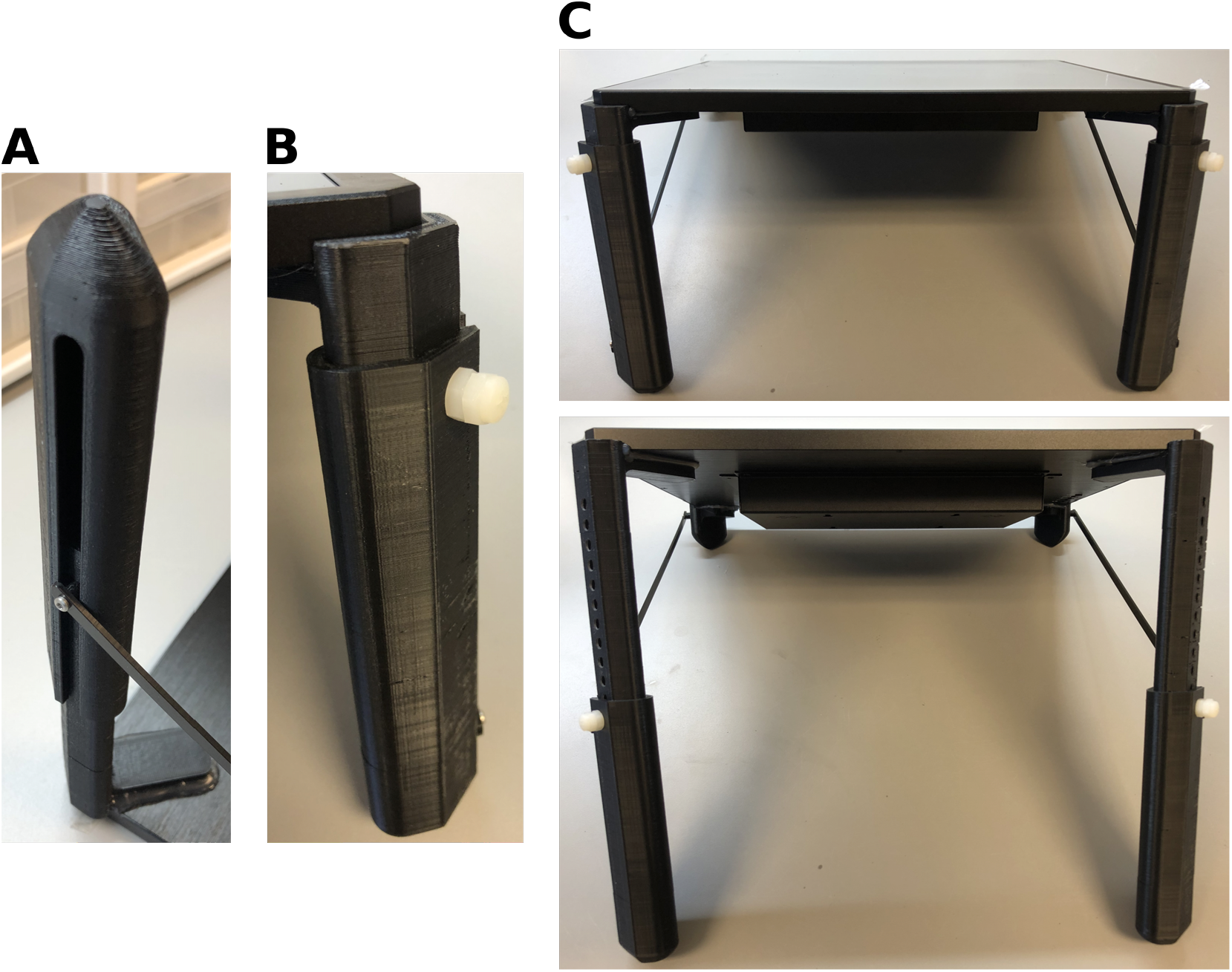
Installation of the leg extensions. (A) The slot on each leg extension is meant to fit around the brace standoff on its respective leg. (B) The nylon screws are used to secure the leg extensions in place during use. (C) Leg extensions fully retracted (top) and fully extended (bottom), resulting in screen-to-table angles of ~ 30° and ~ 46°, respectively.

**Figure 11:**
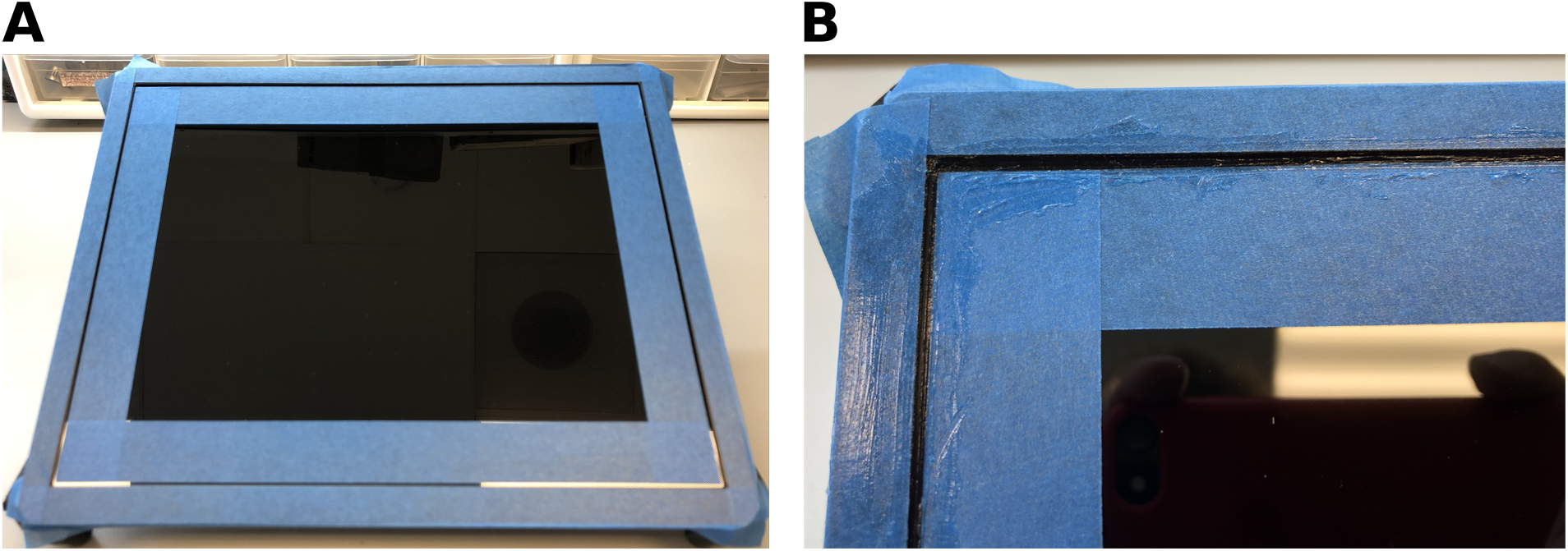
(A) Masking tape applied to the bezel and edges of the screen, leaving a 1-2 mm gap between the edge of the screen and the bezel. (B) Detail of the silicone sealant applied all around the edge of the screen, to seal the gap between the screen and the bezel.

### 5.2 Front panel

1. Parts required
  - Front panel (3-0442 or 3-0443) - laser cut
  - Plate holder(s) (3-0443, 3-0498, or 3-0484) - 3D printed
  - Plate holder screws
  - Panel screws
2. Countersink the back side of the front panel holes used to attach the plate holder (Figure 12A). The countersinks must be deep enough to keep the flat head of the plate holder screws flush or below the surface of the panel. If using M3 screws (instead of #2) the holes on the front panel might have to be drilled to provide proper clearance.
3. Secure the plate holder to the front panel using the plate holder screws (Figure 12B). Make sure the heads of the screws do NOT protrude over the surface of the panel - if they protrude, make the countersinks deeper.
4. Tap the five holes around the periphery of the panel with a #4-40 tap (or M3).
5. Screw the panel screws (#4-40 or M3) all the way into all five holes around the periphery of the panel, with the heads of the screws on the side of the panel where the plate holder is (front side), and tighten lightly. Completed front panels are shown in Figure 13.
6. The build process for the “Well-Lit to Well-Lit” front panel is identical, with the exception of having to mount a second plate holder.
7. The front panel is placed on top of the screen, with the panel screws sliding snugly around the edges of the screen. This arrangement makes it very easy to remove the front panel for cleaning, to replace it with a different panel for a different type of plate, or to alternate between a “Tube to Well-Lit” and a “Well-Lit to Well-Lit” configuration.

**Figure 12:**
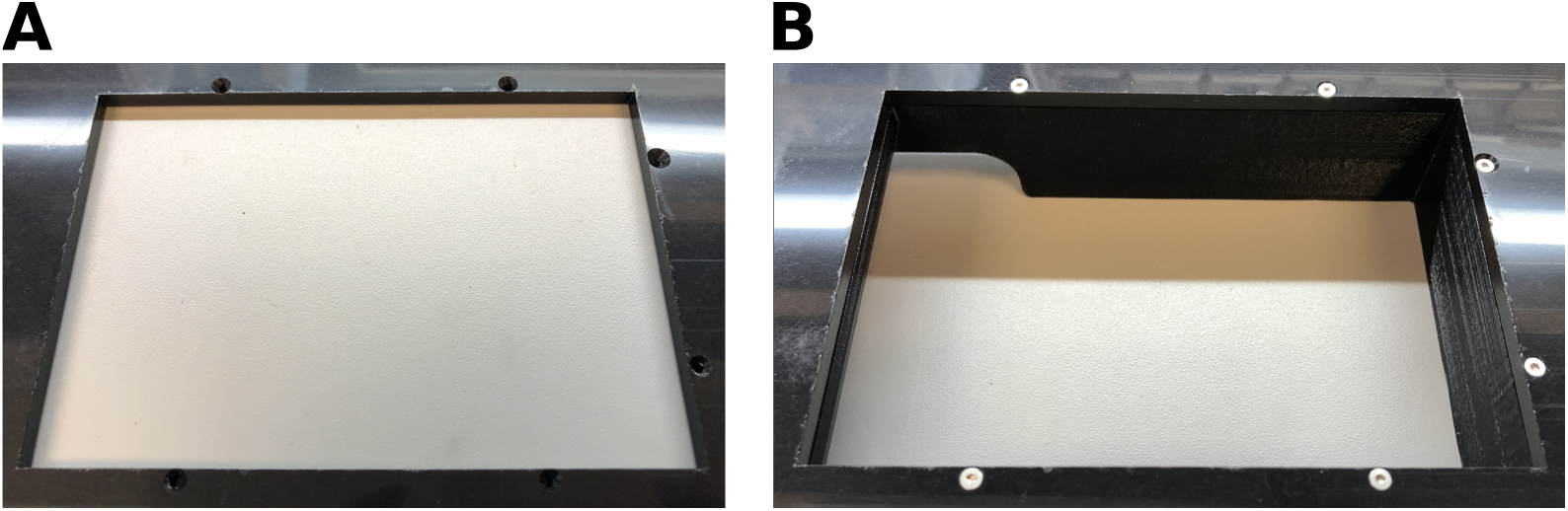
Mounting the plate holder to the front panel. (A) Countersunk holes for the plate holder screws (on the back of the front panel). (B) Plate holder screws tightened into their respective holes on the plate holder. Note that the screw heads are slightly below the surface of the panel.

**Figure 13:**
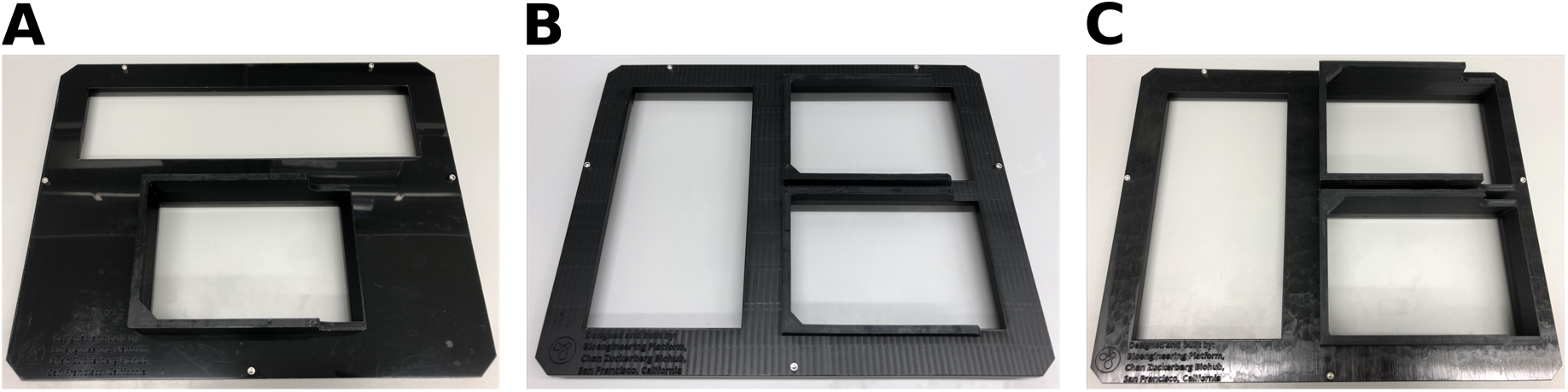
Finished front panels. (A) Front panel for a “Tube to Well-Lit” using the holder for a Fisher Sci. 2 mL deep-well plate (part# 3-0443). (B) Front panel for the “Well-Lit to Well-Lit” for Bio-Rad low profile PCR plates (holder part# 3-0484). (C) Front panel for the “Well-Lit to Well-Lit” for USA Scientific 1 and 2 mL deep-well plates (holder part# 3-0493) (C).

### 5.3 Wiring

1. Use the USB cable provided with the monitor to connect the USB port on the monitor to the mini-PC. This will enable touch screen capabilities.
2. Connect both the mini-PC and the monitor to power.
3. Connect the mini-PC and monitor with the HDMI cable provided with the monitor.
4. For the Tube to Well-Lit version of the device, connect the barcode scanner to a USB port on the mini-PC.

## 6. Installation and configuration

### 6.1 Barcode scanner configuration

This subsection applies only to the Tube to Well-Lit version of the device. If using the same barcode scanner as listed in the bill of materials, users should configure it as follows: Download and print the ‘Well-Lit Barcode Scanner Configuration Sheet.pdf’ file and scan the barcodes on the sheet in a zig-zag pattern, from top to bottom, following the order of the numbers in red. Each barcode in this sheet configures a different aspect of the scanner.

### 6.2 Software installation

The following software installation instructions must be carried out in the computer that will be running the Well-Lit device. As specified in the bill of materials, we have always used a Windows PC for this purpose, but the instructions below should also work on MacOS and Linux, although we have not tested the software in those operating systems. Both versions of the Well-Lit require setting up a Python software environment with several dependencies. The steps are as follows:

- Install Anaconda or Miniconda Python 3.7 (from www.anaconda.com - tested on Anaconda version 4.8.3) selecting the option ‘ADD TO PATH’ in the installer
- Create a new Anaconda environment, activate it, then install dependencies. To do this, open an anaconda command line prompt and enter the following sequence of commands:

~~~
conda create -n WellLit python=3.7.6
conda activate WellLit
conda install matplotlib==3.1.3
conda install -c conda-forge kivy
pip install kivy-garden
garden install graph
garden install matplotlib
~~~
- Clone one or both of the following repositories, depending on the application: https://github.com/czbiohub/WellLit-WelltoWell.git or https://github.com/czbiohub/WellLit-TubeToWell.git
- Open a git bash terminal in each repository folder you just downloaded and enter the command ‘`git submodule update --init`’ to finish obtaining the required files. If you are not using git and downloaded a zipped folder, then download and extract the repository from https://github.com/czbiohub/WellLit.git into the ‘../WellLit’ folder in the first repository you downloaded.

#### 6.2.1 Software configuration

To configure the WellLit software for your device, open ‘wellLitConfig.json’ in a text editor and modify the following entries to suit the application. If invalid directory locations are given in this configuration file, the software will default to using subfolders named ‘samples’, ‘records’, and ‘protocols’ in the parent repository folder.

- `num_wells` configures the software for either 96 or 384 well format. If an invalid number is entered the software defaults to 96-well format.
- `records_dir` configures the directory for storing records. The software automatically records every transfer in a CSV file with timestamps as soon as the action is completed.
- `A1_X_dest` and `A1_Y_dest` control the position of well A1 on the screen. The numeric values are given as fractions of the screen area, and so will likely need to be adjusted if using a screen different than the one specified in this build guide. These values increment from the upper left corner of the Graphical User Interface (GUI). If the lighting pattern is misaligned with the wells on your screen, adjust these parameters to achieve good alignment.
- `size_param` controls the size of the illuminated circle or square which appears beneath a well.
- `well_spacing` controls the distance between adjacent wells.
- *“Tube to Well-Lit” only* : `samples_dir` sets the directory to load CSV files from if the user wishes to restrict plated samples to a pre-defined list.
- *“Tube to Well-Lit” only* : `controls` specifies wells that will be excluded from the sample transfer. If no controls are used this field should be left as empty quotation marks.
- *“Well-Lit to Well-Lit” only* : `protocols_dir` sets the directory to load cherry picking lists. Examples of cherry-picking lists are given in the ‘/protocols’ directory in the repository.
- *“Well-Lit to Well-Lit” only* : `A1_X_dest` and `A1_Y_source` control the position of well A1 for the plate on the bottom half of the screen where samples are aliquoted to.

Both software repositories come with startup launchers that can be copied to the desktop to avoid using a python command line. To use the startup icon, copy ‘startup.bat’ to the desktop and double click it to launch the Well-Lit software.

## 7. Operation instructions

This device is in active use at the Biohub and the software in the public repository will be updated over time to fix bugs and add features. Consequently, the following instructions are only guaranteed to be accurate as of the time of writing/publishing.

### 7.1 Tube to Well-Lit

The “Tube to Well-Lit” version of the device is shown in Figure 14. The top area of the screen contains the main user interface, while the bottom displays the colored dots illuminating the wells. The Graphical User Interface (GUI) will always display the user’s next step for using the device. The GUI will also display error messages if the user attempts to perform invalid commands.

**Figure 14:**
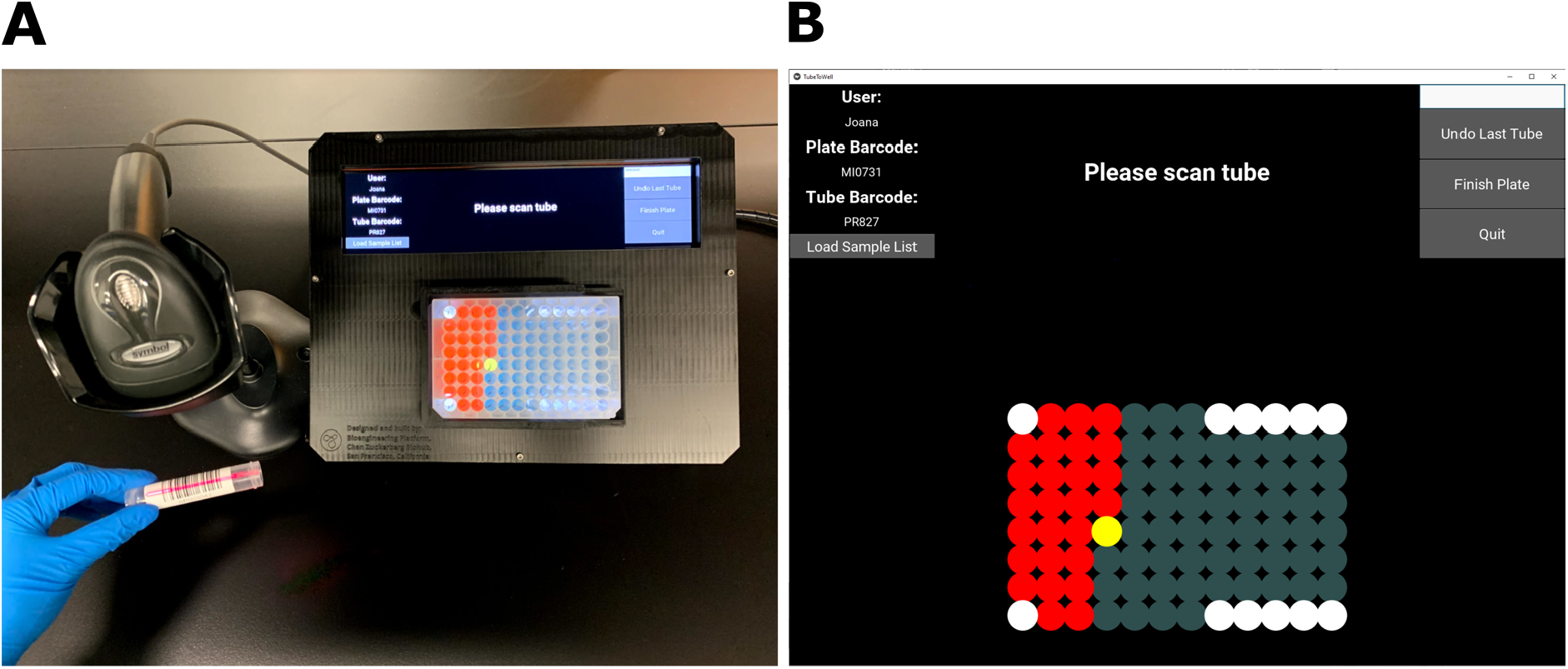
“Tube to Well-Lit” in operation. (A) The user scans the barcode on a tube and the Well-Lit indicates the target well for the contents of the tube in yellow. The blue hue on the buttons and wells in this image is an artifact produced by the automatic white balancing of the camera used to take the photo. (B) GUI of the “Tube to Well-Lit”. The top-left section of the screen displays the user name, plate barcode, the last scanned tube barcode, and the “Load Sample List” button. The top-center section displays instructions for the user. The top-right section has a text entry box (white) and three buttons: “Undo Last Tube”, “Finish Plate”, and “Quit”. The colors in this image accurately represent what the user will see when using the device.

1. Connect the barcode scanner to the mini-PC. The scanner must be configured as described in section 6.1.
2. Ensure that the ‘WellLit-TubeToWell’ repository is active, and that the software configurations have been set.
3. Launch the Well-lit software by double clicking on the ‘startup.bat’ icon, or by launching ‘TubeToWellGUI.py’ from a python terminal.
4. The user will be prompted to enter the user name and the plate name/barcode. All prompted information can either be scanned or entered manually by clicking in the white text entry box on the top-right corner of the screen.
5. Insert the plate into the holder as shown in Figure 14A. Ensure that the A1 well is in the top left corner of the holder (the holder for each type of multi-well plate is designed to ensure that the plate can only be inserted in the right orientation).
6. If the user wishes to restrict tube barcodes to come from a pre-defined list, for example to guard against errors when manually typing by hand or segregating tubes by batches that may have been mixed up, press “Load Sample List” to select a CSV file of tube barcodes. Only barcodes from this list will be accepted by the machine for assigning to a well.
7. Each user action is recorded with a timestamp in a CSV file saved to the folder specified in the the ‘wellLitConfig.json’ configuration file (`records_dir` parameter - see Software Configuration section).
8. Wells are highlighted with the following colors:
  - Yellow: Current transfer target well
  - Red: Full wells
  - Gray: Empty wells
  - Blue: Re-scanned sample well (full)
  - White: Excluded/control wells (see software configuration section)
9. “Tube to Well-Lit” sample transfer procedure:
  - Scan or type a tube name or barcode to light up the first available well in yellow as shown in Figure 14. This is the target well. Wells are assigned to the tubes in a column-wise order (i.e. A1, B1, C1, … A2, B2, C2, …).
  - After transferring an aliquot from the tube to the target (yellow) well, scan or enter the next tube’s barcode to mark the previous transfer as complete and light up the next available well. The filled wells will be lit in red.
  - Re-scanning a previously scanned tube will light its assigned well in blue.
  - The last transfer can be undone with the “Undo Last Tube” button. The undone transfer is not included in the record file generated. You cannot undo more than one transfer per scan.
10. Press “Finish Plate” when all the transfers have been completed. The program will automatically start a new record file for the next plate. For a new plate follow the instructions starting at step 4 for the new plate.

### 7.2 Well-Lit to Well-Lit

The assembled “Well-Lit to Well-Lit” version of the device is shown in Figure 15A. The GUI shown in 15B will display instructions for the user as they use the device. This GUI will also show popups if the user attempts any invalid commands.

**Figure 15:**
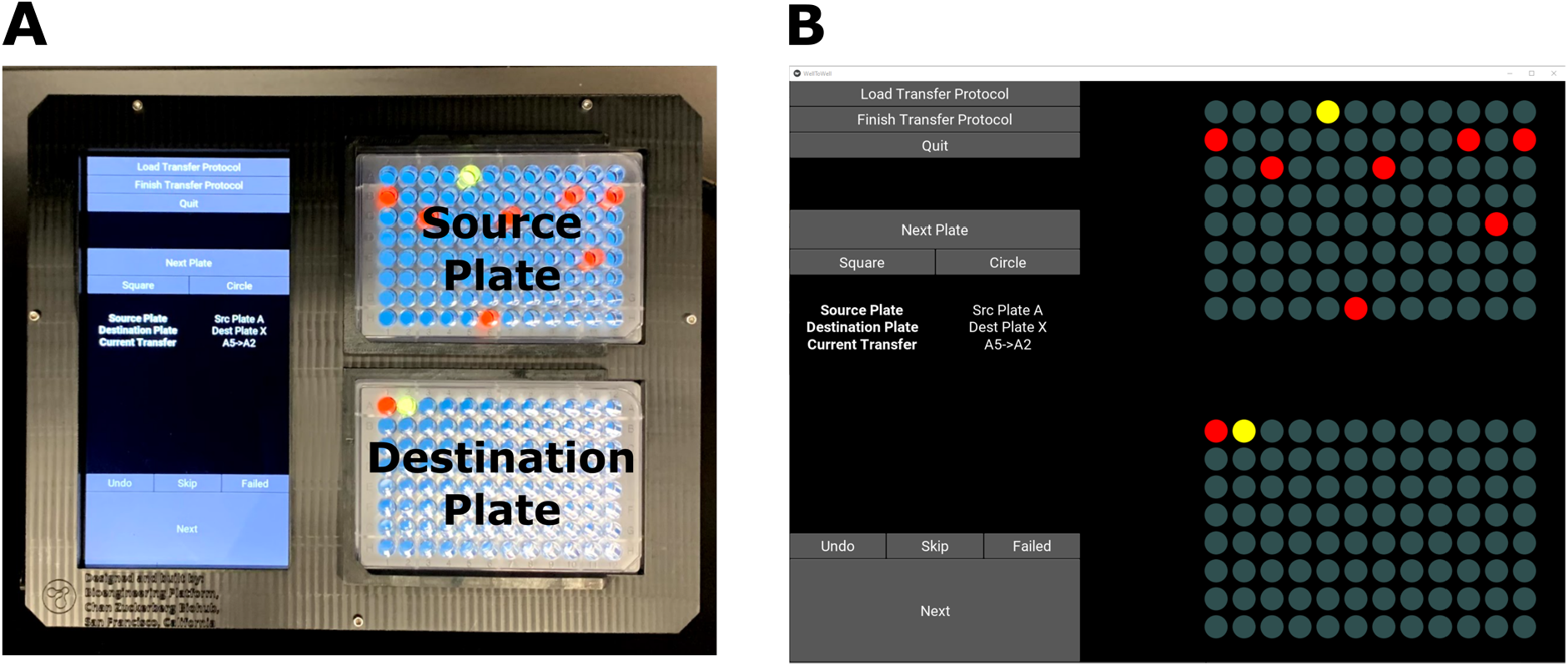
“Well-Lit to Well-Lit” in operation. (A) Two deep well plates mounted onto the device, with a well-to-well transfer in progress. Top plate is always the source; bottom plate is always the destination. The blue hue on the buttons and wells in this image is an artifact produced by the automatic white balancing of the camera used to take the photo. (B) GUI of the “Well-Lit to Well-Lit”. The left section contains all the buttons that the user needs to interact with the device and information about the transfer protocol being run. The colors in this image accurately represent what the user will see when using the device.

1. Ensure that the ‘WellLit-WelltoWell’ repository is active, and that the software configurations have been set.
2. To produce a cherry picking transfer protocol, create a new CSV file in the folder specified in the ‘wellLitConfig.json’ configuration file (`protocol_dir` parameter - see Software Configuration section). Examples of cherry picking CSV files can be seen in the ‘/protocols’ folder in the software repository. The format must be as follows:
  - First Line: Name of destination plate
  - Each line 2…N: Source plate name, source well, destination well
3. Launch the Well-Lit GUI either by double clicking on the ‘startup.bat’ you copied to the desktop, or by launching ‘WellToWellGUI.py’ from a python terminal.
4. Insert the plates into the holders as shown in Figure 15A. Ensure that the A1 wells are in the top left corner of the holder (the holder for each type of multi-well plate is designed to ensure that the plate can only be inserted in the right orientation).
  - Top plate is always the source plate
  - Bottom plate is always the destination plate
5. Load a transfer protocol by clicking on the “Load Protocol” button. The source and destination plate areas will be populated with lights corresponding to each well in the plate. Wells are highlighted with the following colors:
  - Yellow: Source and destination wells for the current transfer
  - Red: Wells that are listed for transfer in the protocol file
  - Gray: Wells that were *not* listed for transfer in the protocol file
6. Each user action is recorded with a timestamp in a CSV file saved to the folder specified in the ‘wellLitConfig.json’ configuration file (`records_dir` parameter - see Software Configuration section).
7. “Well-Lit to Well-Lit” sample transfer procedure:
  - Press “Next” or use the hotkey shortcut ‘n’ to light a source well and its corresponding destination well in yellow.
  - Press “Failed” if the transfer was unsuccessful and should be skipped - it will be marked as ‘Failed’ in the log file.
  - Press “Skip” if you do not wish to complete the current transfer that is lit up in yellow - it will be marked as ‘Skipped’ in the log file.
  - After successfully transferring the sample from the source well to the destination well, press “Next” or use the hotkey shortcut ‘n’. The source well will be lit in gray and the destination well will be lit in red to denote that the source has been emptied and the destination has been filled. The next pair of source and transfer wells will be lit in yellow.
  - The last transfer can be undone with the “Undo” button, giving the user the opportunity to redo it. Only the most recent transfer can be undone (not transfers before the last), and a user cannot mark a transfer as undone after they press the “Next Plate” button, even if that transfer was the most recently completed.
  - To complete a destination plate, press “Next Plate” or use the hotkey shortcut ‘p’. If not all transfers on the current destination plate are complete, the user will be asked to confirm the command. If the user confirms, all of the incomplete transfers are marked as ‘Skipped’ in the log file.
8. When the transfer protocol is complete press on “Complete Transfer Protocol” to finish the transfers and allow a new protocol CSV file to be uploaded.

## 8. Validation and characterization

In March 2020, personnel from the UCSF CLIA laboratory were trained on the operation of the “Tube to Well-Lit”, and they successfully used the device to routinely handle SARS-CoV-2 testing samples for approximately six weeks. After those six weeks, liquid handling robots were fully operational and took over the transfer of samples from tubes to multi-well plates. The CLIA lab personnel reported being extremely satisfied with the performance of the “Tube to Well-Lit”, and indicated that it had dramatically increased the tube to well transfer speed and reduced their stress about making mistakes. Two “Tube to Well-Lit” systems were kept in the CLIA lab to be used in case the liquid handling robots were disabled for repair or maintenance.

At the time of writing, two research groups at the Chan Zuckerberg Biohub have successfully used the device for a variety of experiments (one group running three units of the “Tube to Well-Lit” version, and the other group one unit of the “Well-Lit to Well-Lit” version), and they report that the device has been incredibly helpful in both configurations. The “Well-Lit to Well-Lit” unit has been especially important when cherry-picking from and to 396-well plates. With so many wells, it’s almost impossible to perform cherry-picking transfers without errors in a reasonable amount of time. Both groups report that the device also drastically reduces operator fatigue and stress, compared to unaided manual transfers.

## 9. Acknowledgements

The authors would like to thank Hanna Retallack for thoroughly testing the device under real-use conditions and providing very valuable feedback, Joe DeRisi for encouraging us to build the device in record time, and Lillian Cohn and Cristina Tato and her teams for providing very valuable feedback on the “Well-Lit to Well-Lit” and “Tube to Well-Lit” devices, respectively. This work was supported by the Chan Zuckerberg Biohub.

## 11. Human and animal rights

This work did not involve any human or animal experimental subjects.

